# LMO3 is a suppressor of the basal-like/squamous PDAC subtype and reduces disease aggressiveness of pancreatic cancer through glycerol 3-phosphate metabolism

**DOI:** 10.1101/2023.11.01.564448

**Authors:** Yuuki Ohara, Amanda J. Craig, Huaitian Liu, Shouhui Yang, Paloma Moreno, Tiffany H. Dorsey, Helen Cawley, Azadeh Azizian, Jochen Gaedcke, Michael Ghadimi, Nader Hanna, Stefan Ambs, S. Perwez Hussain

**Author notes:** Address Correspondence to: S. Perwez Hussain, Ph.D., Chief, Pancreatic Cancer Section Laboratory of Human Carcinogenesis, Bldg. 37 Room 3044B, National Cancer Institute, NIH, 37 Convent Dr., Bethesda, MD 20892, Yuuki Ohara, M.D., Ph.D., Bldg. 37 Room 3054, National Cancer Institute, NIH,**/**.

## Abstract

Pancreatic ductal adenocarcinoma (PDAC) encompasses diverse molecular subtypes, including the classical/progenitor and basal-like/squamous subtypes, each exhibiting distinct characteristics, with the latter known for its aggressiveness. We employed an integrative approach combining transcriptomic and metabolomic analyses to pinpoint potential genes contributing to the basal-like/squamous subtype differentiation. Applying this approach to our NCI-UMD-German and a validation cohort, we identified LIM Domain Only 3 (LMO3), a transcription co-factor, as a candidate suppressor of the basal-like/squamous subtype. Reduced LMO3 expression was significantly associated with higher pathological grade, advanced disease stage, induction of the basal-like/squamous subtype, and decreased survival among PDAC patients. *In vitro* experiments demonstrated that *LMO3* transgene expression inhibited PDAC cell proliferation and migration/invasion, concurrently downregulating the basal-like/squamous gene signature. Metabolomic analysis of patient tumors and PDAC cells revealed a metabolic program linked to elevated LMO3 expression and the classical/progenitor subtype, characterized by enhanced lipogenesis and suppressed amino acid metabolism. Notably, glycerol 3-phosphate (G3P) levels positively correlated with LMO3 expression and associated with improved patient survival. Furthermore, glycerol-3-phosphate dehydrogenase 1 (GPD1), a crucial enzyme in G3P synthesis, showed upregulation in LMO3-high and classical/progenitor PDAC, suggesting its potential role in mitigating disease aggressiveness. Collectively, our findings suggest that heightened LMO3 expression reduces transcriptomic and metabolomic characteristics indicative of basal-like/squamous tumors with decreased disease aggressiveness in PDAC patients. The observations describe LMO3 as a candidate for diagnostic and therapeutic targeting in PDAC.

**Graphical abstract:** 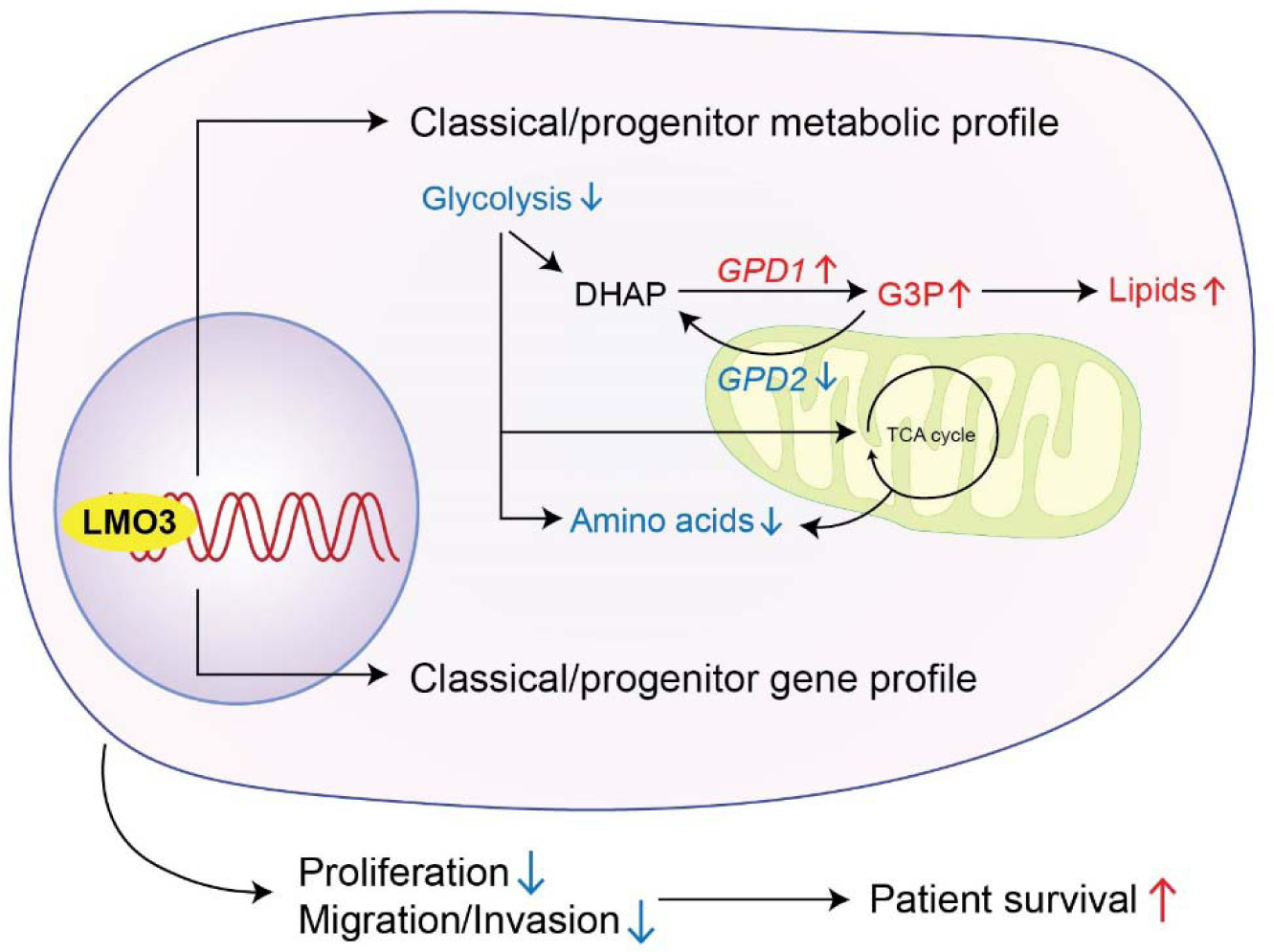

**Highlights:**

- LMO3 is downregulated in basal-like/squamous PDAC while its expression is maintained in the classical/progenitor PDAC subtype
- Upregulated LMO3 expression correlates with improved PDAC survival and reduced proliferation and migration/invasion in PDAC cells
- Upregulated LMO3 suppresses basal-like/squamous differentiation and induces a unique metabolic signature characterized by elevated lipogenesis and diminished amino acid metabolism, resembling the classical/progenitor PDAC subtype
- Enhanced LMO3 expression associates with elevated glycerol 3-phosphate levels in PDAC, correlating with improved patient survival in PDAC

## Introduction

Pancreatic cancer is a highly lethal malignancy, with a meager 5-year survival rate of merely 12%^1^. Accounting for over 95% of pancreatic malignancies, pancreatic ductal adenocarcinoma (PDAC) is its most prevalent form^2^. Within the realm of PDAC research, distinct molecular subtypes have emerged, each characterized by unique biological attributes that hold implications for patient survival^3–5^. *Bailey* et al. introduced a classification system comprising four PDAC subtypes: aberrantly differentiated endocrine exocrine (ADEX), pancreatic progenitor, squamous, and immunogenic^3^. The pancreatic progenitor and ADEX subtypes share the upregulation of transcriptional networks linked to pancreatic development and the differentiation of exocrine and neuroendocrine lineages^3^. The squamous subtype stands out as an aggressive form of PDAC, marked by specific traits such as metabolic reprogramming, inflammation, squamous differentiation, hypoxia, proliferation, and MYC activation^3^. Recent refinements in PDAC classification have led to the categorization of two major subtypes: the ‘classical/progenitor’ and ‘basal-like/squamous’ subtypes’^6^. The ‘classical/progenitor’ subtype maintains a closer connection to normal pancreatic tissue, reflecting an endodermal-pancreatic identity, while the ‘basal-like/squamous’ subtype exhibits traits associated with squamous differentiation^7^.

Metabolic reprogramming, a key characteristic of cancer^8–10^, plays a significant role in PDAC^10–12^. Distinct molecular subtypes of PDAC exhibit varying metabolic profiles, with the classical/progenitor subtype potentially relying more on lipogenesis for its metabolic needs, whereas the basal-like/squamous subtype predominantly utilizes glycolysis^13–16^. However, the intricate relationship between metabolic adaptations and gene expression profiles in PDAC remains insufficiently explored, often constrained by small sample sizes. Building on our previous identification of SERPINB3 as an oncogenic driver of the basal-like/squamous subtype^17^, this study investigated the interplay between gene expression and metabolic reprogramming in the development of the classical/progenitor subtype in PDAC. Our results highlight the role of LIM Domain Only 3 (LMO3) upregulation in shaping the metabolic landscape of the classical/progenitor subtype in PDAC.

## Materials and Methods

### PDAC cohorts

We employed gene expression datasets from the “Bailey” cohort (GSE36924)^3^, the “Moffitt” cohort (GSE71729)^4^, and our NCI-UMD-German cohort (GSE183795)^18^ to identify genes associated with the development of PDAC molecular subtypes. Comparative analysis between the basal-like/squamous and classical/progenitor subtypes was performed using Partek Genomics Suite 7.0 (Partek Inc., Chesterfield, MO). For the integrated transcriptomic and metabolomic investigations, we utilized RNA sequencing data from patient PDAC tumors within the NCI-UMD-German cohort (GSE224564)^17^.

### Cell lines and culture condition

Human PDAC cell lines were purchased from American Type Culture Collection (ATCC, Rockville, Maryland). Authentication of all cell lines via short tandem repeat (STR) profiling had been conducted within the past three years. All experiments were performed using mycoplasma-free cells. For CFPAC-1 (RRID:CVCL_1119) and Capan-1 (RRID:CVCL_0237) cell lines, IMDM supplemented with 10% FBS and 1% penicillin-streptomycin was used. The Capan-2 (RRID:CVCL_0026) cell line was cultured in McCoy’s 5A (Modified), also supplemented with 10% FBS and 1% penicillin-streptomycin. For the remaining PDAC cell lines (AsPC-1; RRID:CVCL_0152, BxPC-3; RRID:CVCL_0186, MIA PaCa-2; RRID:CVCL_0428, PANC-1; RRID:CVCL_0480, Panc 10.05; RRID:CVCL_1639, SU.86.86; RRID:CVCL_3881), RPMI 1640 with GlutaMax^TM^, 10% FBS, and 1% penicillin-streptomycin was used. All cell cultures were maintained in a humidified incubator with 5% CO2 at 37°C, with all cell culture reagents procured from Thermo Fisher Scientific (Waltham, MA).

### LMO3 overexpression after lentiviral infection

To establish stable cell lines overexpressing *LMO3*, the *LMO3* construct (EX-Z0413-Lv103) and the corresponding empty vector control (EX-NEG-Lv103), both sourced from Genecopoeia (Rockville, MD), were employed. Lentiviral particles were generated by transfecting 293T cells with the lentiviral expression vectors and the Lenti-Pac^TM^ HIV Expression Packaging system, also from Genecopoeia. To obtain stable clones of PDAC cells (Panc 10.05 and SU.86.86), selection was carried out using 4 μg/ml puromycin (Thermo Fisher Scientific).

### RNA sequencing

Quadruplicate RNA sequencing was conducted using total RNA isolated from the human PDAC cell line (Panc 10.05 +/− LMO3). PDAC cells were cultured in RPMI 1640 medium, GlutaMaxTM, supplemented with 10% FBS and 1% penicillin–streptomycin for 72 hours prior to RNA extraction. Libraries were prepared by the Sequencing Facility at NCI-Leidos using the TruSeq Stranded mRNA Kit (Illumina, San Diego, CA) and sequenced in a paired-end manner on NextSeq (Illumina) with 2 x 101 bp read lengths. In summary, this generated approximately 32 to 60 million paired-end reads with a base call quality of ≥ Q30. The fastq-format sequence reads were then aligned to the human reference genome hg38 using STAR and RSEM to obtain gene expression data, reported as transcripts per million with FPKM mapped reads. Differential expression analysis was conducted using DESeq2. Enrichment analysis covering established pathways and datasets was performed using Ingenuity pathway analysis (IPA, QIAGEN, Venlo, Netherlands) and Gene Set Enrichment Analysis (GSEA). The RNA sequencing data have been deposited in the NCBI’s Gene Expression Omnibus (GEO) database under accession number GSE246660.

### Quantitative real-time PCR for *LMO3*

The High-Capacity cDNA Reverse Transcription Kit (Thermo Fisher Scientific) was employed to synthesize first-strand cDNA from total RNA. Quantitative real-time PCR (qPCR) assays were conducted using Taqman probes (Thermo Fisher Scientific) for *LMO3* (Hs00998696_m1) and *GAPDH* (Hs99999905_m1).

### Metabolic profiling and data analysis of PDAC

Tumor sample metabolic profiling was conducted by Metabolon Inc. (Morrisville, NC) following their standard protocol^19–22^. Metabolon’s untargeted metabolic platform utilizes two separate ultra-high performance liquid chromatography/tandem mass spectrometry (UHPLC/MS/MS) injections and one gas chromatography/mass spectrometry injection for each sample, enabling the comprehensive measurement of all metabolites. The resulting dataset from the NCI-UMD-German cohort (n = 50) was normalized across samples before delivery, in accordance with Metabolon’s established protocol, and has been archived in a repository^23^. The normalized relative abundance levels for each metabolite were collated and employed for subsequent data analysis. The same standardized protocol was applied to perform metabolic profiling of cultured cells. Here, we conducted quadruplicate metabolic profiling of the human PDAC cell line (Panc 10.05 +/− LMO3). The PDAC cells were cultivated in RPMI 1640 with GlutaMax^TM^, supplemented with 10% FBS and 1% penicillin–streptomycin for 72 hours. Subsequently, cell pellets were collected, preserved at −80 °C, and sent to Metabolon Inc. for analysis. For enrichment analysis, we utilized MetaboAnalyst 5.0 (https://www.metaboanalyst.ca).

### Cell proliferation assay

The CCK-8/WST-8 assay was utilized to assess the proliferation of PDAC cells (Panc 10.05 and SU.86.86). We seeded 5×10^3^ cells in a 96-well plate with RPMI 1640 supplemented with GlutaMax^TM^, either 10% FBS or 1% FBS, and 1% penicillin-streptomycin. The proliferation assay was conducted at 24, 48, 72, and 96 hours post-seeding, following the manufacturer’s protocol (Dojindo Laboratories, Kumamoto, Japan). Absorbance readings were obtained using a SpectraMax® ABS Plus microplate reader (Molecular Devices, San Jose, CA).

### Cell migration and invasion assay

The migration assay was conducted with 24-well Falcon^®^ Cell Culture Insert (Corning, Glendale, AZ). For invasion assay, Matrigel Basement Membrane Matrix (#354234, Corning) was used. The upper chamber’s membrane was coated with 100 μl of Matrigel matrix coating solution (Matrigel matrix: coating buffer = 1:39) for 2 hours, adhering to the manufacturer’s guidelines. The lower chamber was filled with 750 μl of RPMI 1640, GlutaMax^TM^ supplemented with 10% FBS, while the upper chamber was loaded with PDAC cells (Panc 10.05; 10×10^4^ cells, SU.86.86 5×10^4^ cells) in 500 μl of serum-free RPMI 1640, GlutaMax^TM^. Cells were then incubated for 48 hours in a humidified incubator at 5% CO2 and 37°C. After incubation, cells that had migrated or invaded through the membrane were fixed using 100% methanol (Thermo Fisher Scientific) and subsequently stained with Crystal violet solution (MilliporeSigma, Burlington, MA). Finally, the cells were counted.

### Immunohistochemistry

Four μm thick paraffin-embedded tumor sections, as well as adjacent non-tumor sections, were subjected to overnight incubation at 4°C with a rabbit monoclonal anti-LMO3 antibody (Abcam, Cambridge, UK, ab230490, 1:500). Signal amplification was achieved using the Dako envision+ system-HRP labeled polymer anti-rabbit antibody (Agilent Technologies, Santa Clara, CA), followed by color development with 3,3’-diaminobenzidine (DAB, Agilent Technologies). Immunohistochemistry (IHC) was assessed by assigning intensity and prevalence scores^17, 22^. Intensity scores ranging from 0 to 3, representing negative, weak, moderate, or strong expression, were assigned. Prevalence scores ranging from 0 to 4, representing the percentage of cells showing LMO3 expression (<10%, 10–30%, >30–50%, >50–80%, and >80%), were also assigned. The overall IHC score was obtained by multiplying the intensity and prevalence scores.

### Immunoblotting

PDAC cells were lysed using RIPA Lysis and Extraction Buffer from Thermo Fisher Scientific, supplemented with cOmplete™ Protease Inhibitor Cocktail (MilliporeSigma, 4693116001). The protein extracts were subjected to electrophoresis under reducing conditions on 4–15% polyacrylamide gels (Bio-Rad Laboratories, Inc., Hercules, CA) and subsequently transferred onto a nitrocellulose membrane (Bio-Rad Laboratories, Inc.). The membrane was blocked with SuperBlock™ Blocking Buffer (Thermo Fisher Scientific) for 1 hour at room temperature, followed by an overnight incubation at 4°C with primary antibodies. Primary antibodies included LMO3 (Abcam, ab230490, 1:200) and β-Actin (MilliporeSigma, A5441, 1:2000). Afterward, the membrane was incubated with secondary ECL anti-rabbit or anti-mouse IgG HRP-linked antibodies (GE Healthcare, Pittsburgh, PA) for 1 hour at room temperature. Protein visualization was achieved using SuperSignal™ West Dura Extended Duration Substrate (Thermo Fisher Scientific).

### Statistical analysis

Statistical analyses were conducted using GraphPad Prizm 9 (GraphPad Software, La Jolla, CA). The difference in overall survival between patient groups was determined using the Kaplan-Meier method and the log-rank test. Group differences were assessed via unpaired two-tailed Student’s t-tests (for two groups) or ANOVA (for three or more groups). Results are presented as mean ± SD, with statistical significance defined as a p-value less than 0.05.

## Results

### LMO3 is downregulated in the basal-like/squamous PDAC and associates with advanced stage and poor patient survival

Our study aimed at identifying potential suppressor genes implicated in the development of the basal-like/squamous subtype of PDAC^23^. Through transcriptome analysis of three PDAC cohorts (Bailey cohort^3^, Moffitt cohort^4^, and NCI-UMD-German cohort^18^), we narrowed down our candidate gene list to 60 genes^23^ (Figure 1A). Among these genes, *LONRF2* and *LMO3* held the highest ranks as the first and second ranked genes, respectively, displaying significantly lower hazard ratios (hazard ratio 0.39; p < 0.05 and 0.40; p < 0.05). Notably, LMO3 exhibited a more substantial downregulation in tumor tissues compared to non-tumor tissues. The protein encoded by *LMO3* acts as a transcriptional co-factor and has previously been identified as a regulator of adipogenesis, which contributes to lipid accumulation^24^. Prior studies, including our own, have demonstrated that lipogenesis can play a critical role in inhibiting the progression of PDAC^25–27^. Building on these observations, we hypothesized that LMO3 could inhibit PDAC progression, or the differentiation into the aggressive basal-like/squamous subtype, through metabolic reprogramming, prompting us to select it for in-depth investigation. Our analysis revealed that reduced *LMO3* transcript levels correlated with decreased patient survival. Additionally, *LMO3* transcripts displayed downregulation in tumor tissues compared to non-tumor tissues in both our NCI-UMD-German cohort and the validation cohort (Moffitt cohort^4^) (Figure 1B-E). Furthermore, the downregulation of LMO3 exhibited a direct association with advanced-stage (III/IV) PDAC and higher-grade tumors (Figure 1F and 1G). Immunohistochemistry (IHC) illustrated the presence of LMO3 protein in the cytoplasm of acinar and endocrine cells, albeit at lower levels in tumor cells (Figure S1A and S1B). These collective findings support the hypothesis that upregulated LMO3 may have a potential inhibitory role in PDAC whereas its diminished expression is associated with adverse patient survival and differentiation into the basal-like/squamous subtype.

**Figure 1.**
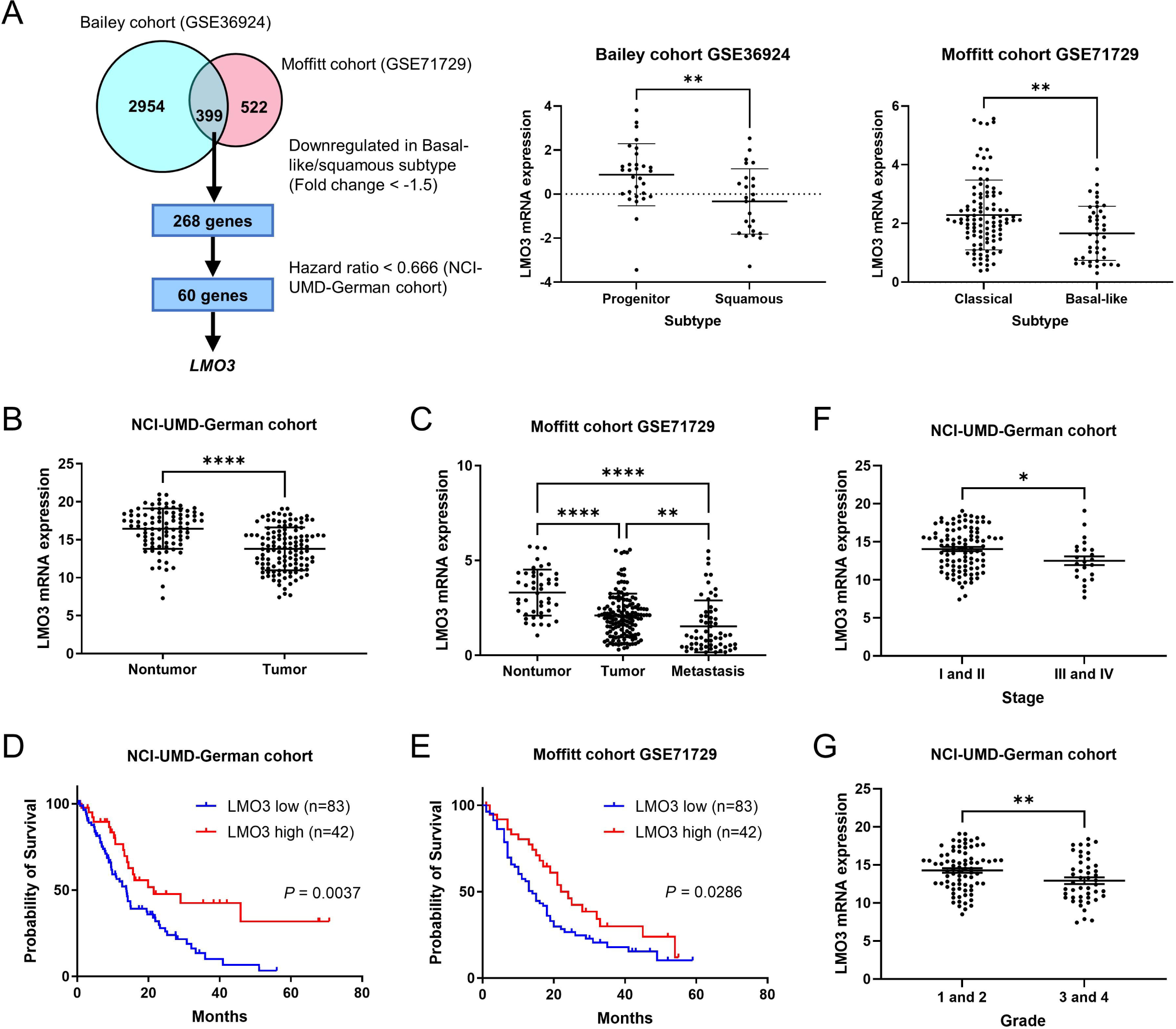
Transcriptome analysis identifies LMO3 as a candidate suppressor of basal-like/squamous PDAC and a prognostic marker of patient survival. (A) Workflow for the identification of candidate suppressor genes in the basal-like/squamous subtype. Candidate genes were identified through the analysis of two cohorts (Bailey cohort^3^ and Moffitt cohort^4^), followed by survival analysis in the NCI-UMD-German cohort^18^. (B-E) Comparison of *LMO3* transcript levels in PDAC tumors and adjacent non-cancerous tissues (B and C). *LMO3* expression is downregulated in tumors, as observed in both the NCI-UMD-German cohort (qPCR) and a validation cohort (Moffitt cohort^4^). Metastatic PDAC shows the lowest LMO3 expression (C). Panels D and E show Kaplan-Meier plots and log-rank test results, depicting the association between decreased *LMO3* and decreased PDAC patient survival in the NCI-UMD-German and a validation cohort (Moffitt cohort^4^). The comparison was made between patients in the upper 33.3% tertile and the lower 66.6% of *LMO3* expression. (F-G) Downregulation of *LMO3* is evident in advanced-stage (III/IV) and higher-grade PDAC (qPCR). Data represent mean ± SD. *p ≤ 0.05, **p ≤ 0.01, ***p ≤ 0.001, ****p ≤ 0.0001 by unpaired two-tailed Student’s t-test or one-way ANOVA.

### Upregulation of LMO3 downregulates proliferation and cellular movement in PDAC

We next investigated whether LMO3 expression would mitigate the characteristics associated with the basal-like/squamous subtype, thereby reducing disease aggressiveness. In a prior study, we characterized the molecular subtypes of 175 patients in our NCI-UMD-German cohort (unclassified; n = 32, classical/progenitor; n = 95, basal-like/squamous; n = 48)^17^. We showed that pathways related to cellular movement exhibited heightened activity in the basal-like/squamous subtype compared to the classical/progenitor subtype. Extending this work, we found that tumors categorized as the basal-like/squamous subtype displayed lower levels of *LMO3* mRNA expression compared to other subtypes (Figures S2A). Pathway enrichment analysis using the Ingenuity Pathways Analysis (IPA) unveiled that LMO3-high PDAC tumors enriched the pathways related to cellular movement and cellular growth (Figure 2A). To validate these observations, we assessed LMO3 expression in human PDAC cell lines, revealing that LMO3 expression remained consistently low to undetectable across all the PDAC cell lines (Figure S2B and S2C). We selected Panc 10.05 and SU.86.86 cells to establish cell lines with *LMO3* transgene expression. The presence of upregulated LMO3 expression was confirmed at both the mRNA and protein levels in these cell lines (Figure S2D and S2E). IPA revealed concordance among enriched pathways in PDAC cells with *LMO3* transgene expression and LMO3-high PDAC tumors in the NCI-UMD-German cohort, highlighting the significance of pathways associated with cellular movement and growth (Figure 2B). To further investigate LMO3’s functional role, we performed additional *in vitro* experiments using these LMO3-overexpressing PDAC cells. The CCK-8/WST-8 assay provided evidence that LMO3 expression impeded proliferation, with this inhibitory effect becoming even more apparent under conditions of limited nutrient supply (Figure 2C). Furthermore, LMO3 expression significantly inhibited the migration and invasion of PDAC cells (Figure 2D). Additionally, our analyses employing gene set enrichment analysis (GSEA) revealed that LMO3 upregulated neuroendocrine differentiation (Figure 3A and 3B), a characteristics feature associated with the classical/progenitor subtype^3^. Simultaneously, it downregulated the basal-like gene signature^28^ (Figure 3A and 3B, right panel). Collectively, these findings suggest that LMO3 plays a role in suppressing the characteristics linked to the basal-like/squamous subtype of PDAC.

**Figure 2.**
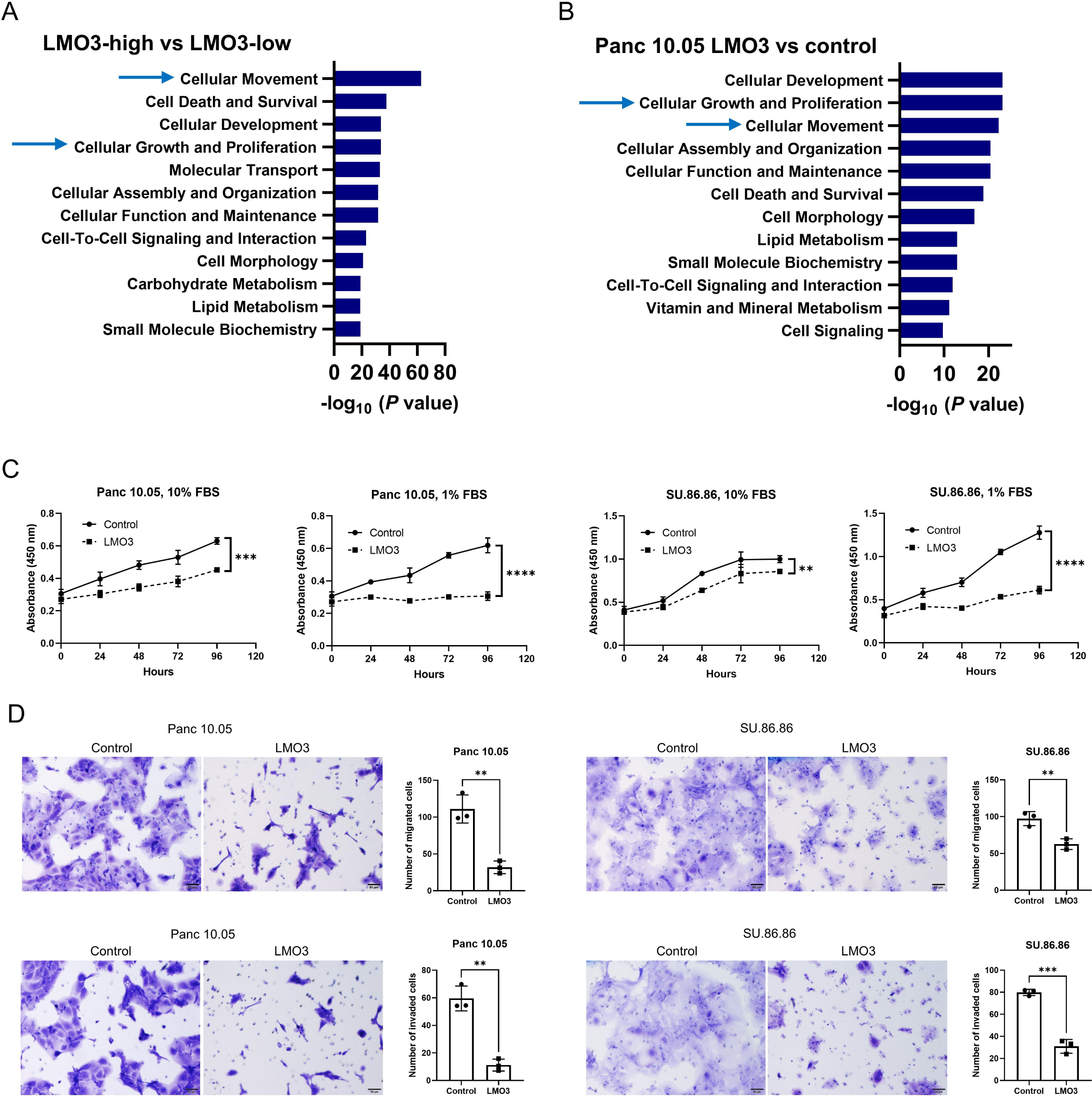
LMO3 suppresses proliferation, migration, and invasion in PDAC. (A-B) Transcriptomic analyses of patient PDAC and Panc 10.05 human PDAC cells. Ingenuity pathway analysis (IPA) of contrasts LMO3-high versus LMO3-low PDAC tumors and Panc 10.05 cells with *LMO3* transgene expression (LMO3-high) versus vector control (LMO3-low). IPA enrichment scores (*P* value-based) illustrate pathways enriched by increased LMO3. IPA highlights the association of elevated LMO3 with similar pathways in both patient PDAC and cultured cells, including cellular movement and cellular growth in LMO3-high PDAC tumors and in human PDAC cell lines with *LMO3* transgene overexpression. (C) PDAC cells overexpressing LMO3 exhibit reduced proliferation compared to vector control cells, with this inhibitory effect becoming even more pronounced under conditions of limited nutrient supply (1% FBS). (D) PDAC cells overexpressing LMO3 display diminished migration and invasion capabilities compared to vector control cells. Data represent mean ± SD of three replicates. *p ≤ 0.05, **p ≤ 0.01, ***p ≤ 0.001, ****p ≤ 0.0001 by two-way ANOVA (C) or unpaired two-tailed Student’s t-test (D).

**Figure 3.**
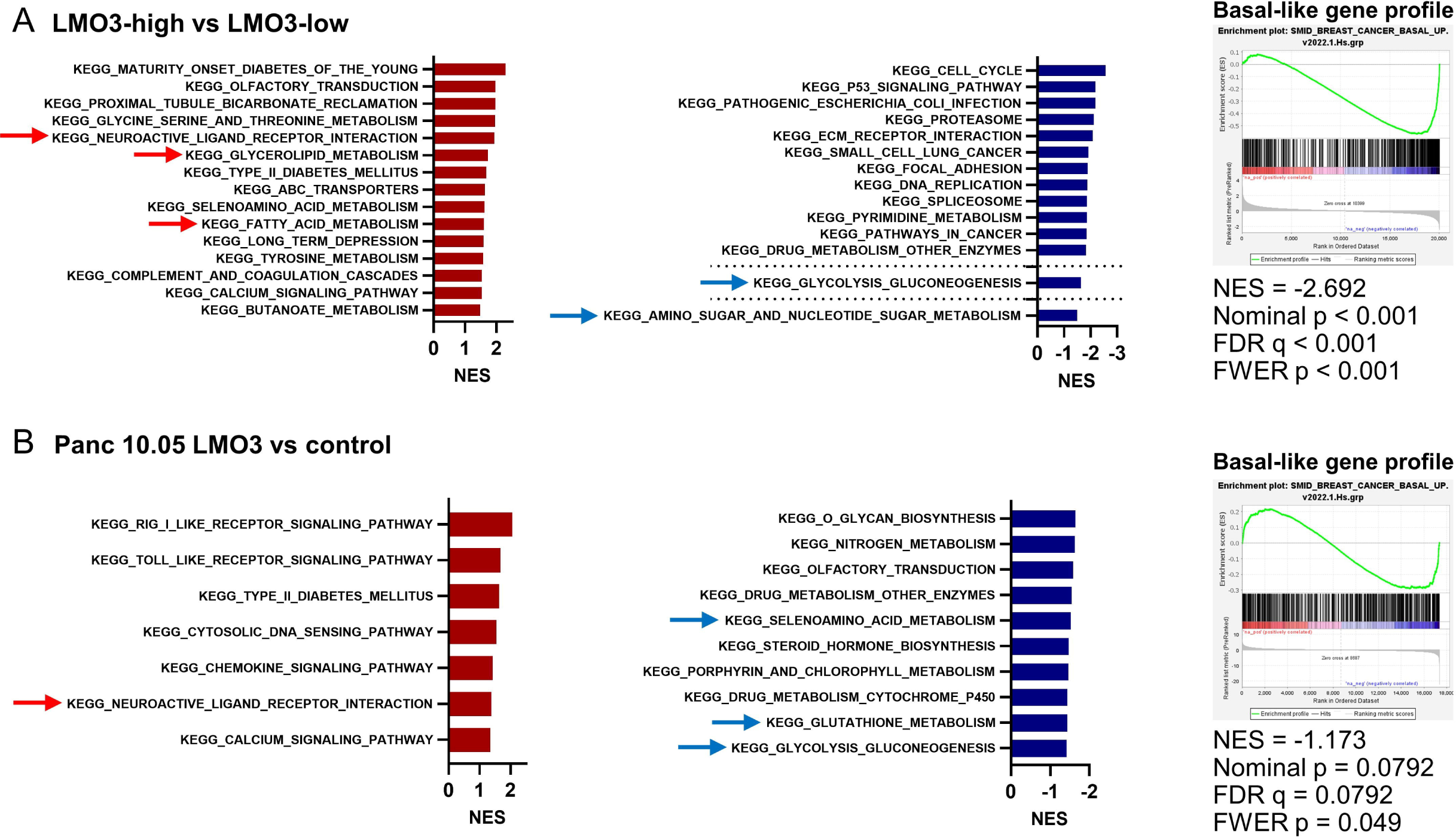
LMO3 upregulation promotes classical/progenitor subtype differentiation and metabolic reprogramming in PDAC. (A-B) Gene Set Enrichment Analysis (GSEA) using transcriptome data from the contrast LMO3-high versus LMO3-low PDAC tumors (A) and the contrast Panc 10.05 cells +/− *LMO3* transgene expression (LMO3-high versus LMO3-low) (B). GSEA suggests that high LMO3 activates neuron differentiation and lipogenesis (left bar graphs), while downregulating pathways associated with glycolysis and amino acid metabolism (middle bar graphs). GSEA also suggests that upregulated LMO3 expression downregulates basal-like/squamous subtype differentiation, as indicated by negative enrichment scores in a predefined basal-like gene profile^28^ from a GSEA dataset (right panels).

### Upregulation of lipid metabolism and downregulation of amino acid metabolism in LMO3-high PDAC

We proceeded to examine the impact of LMO3 on PDAC metabolism. By applying GSEA to the transcriptome data, we observed that LMO3 overexpression led to an upregulation of lipogenesis while concurrently downregulating glycolysis and amino acid metabolism (Figure 3A and 3B).

Expanding our investigation to include metabolomic analysis, we employed transcriptome and metabolome analyses on a cohort of 50 patients from our NCI-UMD-German cohort. We identified 63 significantly upregulated and 69 downregulated metabolites in *LMO3*-high tumors when compared to *LMO3*-low tumors (Figure 4A and Table S1). Pathway enrichment analysis using MetaboAnalyst 5.0 (https://www.metaboanalyst.ca) with these differential metabolites as input highlighted the upregulation of lipid metabolism and the downregulation of amino acid metabolism in LMO3-*high* tumors (Figure 4B and 4C). To delve deeper into the roles of these metabolites, we conducted a comprehensive metabolome analysis in Panc 10.05 PDAC cells. We found that *LMO3* transgene overexpression resulted in the upregulation of 126 metabolites and the downregulation of 134 metabolites when compared to vector control cells (Figure 4D and Table S2). Applying pathway enrichment analysis using MetaboAnalyst 5.0, we found that LMO3-overexpressing Panc 10.05 cells exhibited heightened lipogenesis and diminished amino acid metabolism (Figure 4E and 4F). Our findings align with previous research from our group, which showed that the basal-like/squamous subtype promotes amino acid metabolism whereas classical/progenitor tumors favor increased lipogenesis^17, 23^. Notably, we observed analogous metabolic adaptation patterns in basal-like/squamous PDAC and LMO3-low PDAC cells, potentially contributing to disease progression.

**Figure 4.**
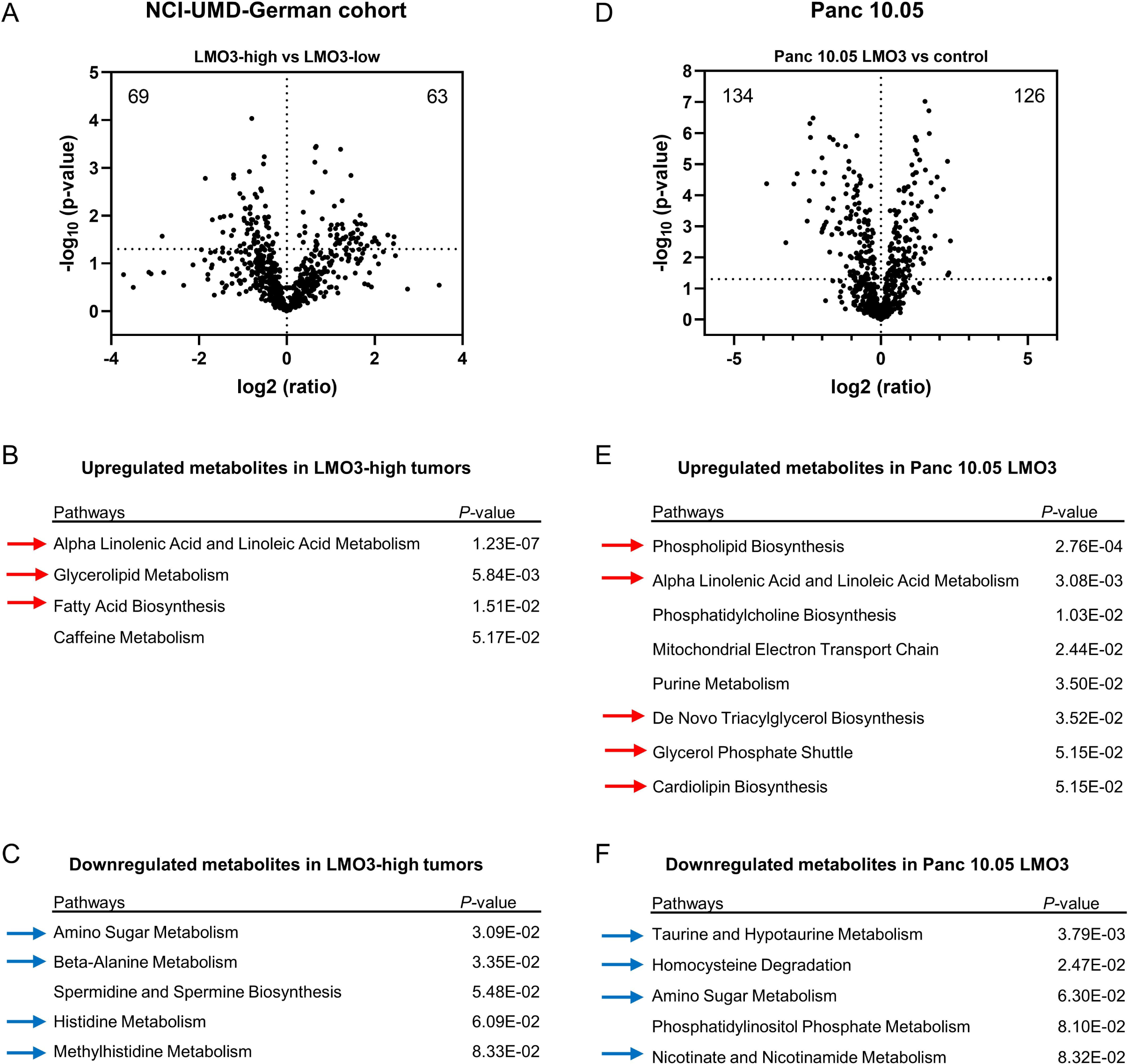
Upregulation of lipid metabolism and downregulation of amino acid metabolism in LMO3-high PDAC. Metabolome analysis of PDAC tumors (NCI-UMD-German cohort) and Panc 10.05 human PDAC cells. Shown are the findings contrasting LMO3-high (n = 25) versus LMO3-low (n = 25) PDAC tumors and Panc 10.05 cells +/− *LMO3* transgene expression (LMO3-high versus LMO3-low). (A) Volcano plot displaying metabolites with differential abundance in LMO3-high versus LMO3-low tumors. 63 metabolites are significantly upregulated and 69 are downregulated in LMO3-high tumors (p < 0.05). The dotted line indicates -log_10_ (p = 0.05). (B-C) Pathway enrichment analysis with differential metabolites using MetaboAnalyst 5.0, highlighting upregulated lipogenesis-related pathways (red arrows) and downregulated amino acid metabolism (blue arrows) in LMO3-high tumors. (D) Volcano plot illustrating metabolites with differential abundance in LMO3-overexpressing versus vector control Panc 10.05 cells. 126 metabolites are significantly upregulated and 134 are downregulated in LMO3-overexpressing Panc 10.05 cells. The dotted line indicates -log_10_ (p = 0.05). (E-F) Pathway enrichment analysis with differential metabolites using MetaboAnalyst 5.0, demonstrating upregulated lipogenesis (red arrows) and downregulated amino acid metabolism (blue arrows) in LMO3-overexpressing Panc 10.05 cells.

### Glycerol 3-phosphate is correlates with favorable patient survival

It was our goal to identify metabolites under the regulation of LMO3 expression. To achieve this, we conducted a metabolome analysis using data from two PDAC datasets, which unveiled 48 metabolites that exhibited significant differential expression between LMO3-high and LMO3-low PDAC (p < 0.05). Among these metabolites, 6 were upregulated, and 21 were downregulated in both datasets (Figure 5A and Table S3). Importantly, all six of the upregulated metabolites fell within the lipid classification. To evaluate the metabolic alterations in the basal-like/squamous subtype, we defined classical/progenitor subtype plus unclassified subtype as non-basal-like/squamous subtype. These 6 metabolites were also elevated in the non-basal-like/squamous subtype within the NCI-UMD-German cohort (Figure S3A). For a more comprehensive analysis, we employed an additional metabolome dataset generated in our previous studies^17, 22^, which comprised a larger patient cohort (a total of 88 patients), albeit with a slightly smaller set of metabolites. Within this dataset, glycerol 3-phosphate (G3P) emerged as a metabolite exhibiting an association with improved patient prognosis (Figure 5B). Notably, glycerol-3-phosphate dehydrogenase 1 (GPD1), an essential enzyme for synthesizing G3P from dihydroxyacetone phosphate (DHAP) and promoting downstream lipogenesis^29–31^, has been linked to anticancer effects^31^. We focused on GPD1 and observed an upregulation of both *GPD1* transcript levels and G3P in LMO3-high PDAC and the non-basal-like/squamous subtype within the NCI-UMD-German cohort (Figure 5C). This trend was also observed in the Bailey cohort^3^ (Figure 5D). In contrast, *GPD2*, which catalyzes the reversible pathway^29, 30^, displayed downregulation in these patient groups (Figure S3B and S3C). Our findings suggest that LMO3-induced metabolic alterations may play a pivotal role in reducing disease aggressiveness in PDAC.

**Figure 5.**
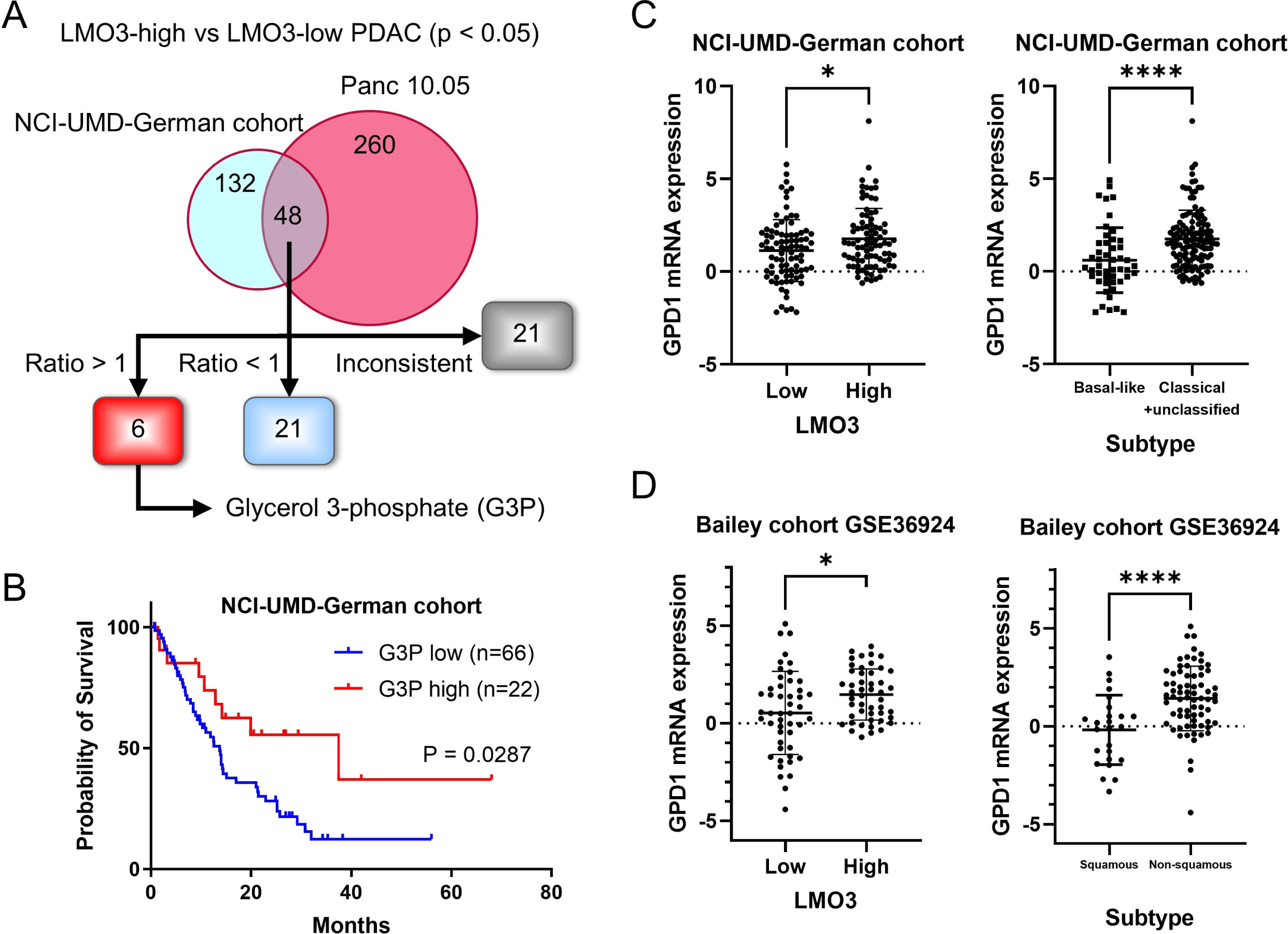
Glycerol 3-phosphate correlates with favorable patient survival in PDAC. (A) Strategy employed to identify metabolites regulated by LMO3, based on analysis of two datasets (NCI-UMD-German cohort^23^ and Panc 10.05 cell line). Forty-eight metabolites exhibited significant differential expression (p < 0.05) in both datasets. Among the identified metabolites, 6 exhibited consistent upregulation (ratio > 1), 21 were consistently downregulated (ratio < 1) in both datasets, while the remaining 21 displayed discordant patterns, being upregulated in one dataset and downregulated in the other. (B) The Kaplan-Meier plot and log-rank test indicate that increased glycerol 3-phosphate (G3P) levels in tumors are associated with improved patient survival (NCI-UMD-German cohort). The comparison was conducted between patients in the upper 25% quartile and the lower 75% of G3P levels. (C) *GPD1* (glycerol-3-phosphate dehydrogenase 1) showed upregulation in both LMO3-high PDAC and the non-basal-like/squamous subtype (classical/progenitor + unclassified PDAC) within the NCI-UMD-German cohort. (D) *GPD1* also displayed upregulation in both LMO3-high PDAC and the non-squamous subtype tumors in the Bailey cohort^3^. Data represent mean ± SD. *p ≤ 0.05, **p ≤ 0.01, ***p ≤ 0.001, ****p ≤ 0.0001 by unpaired two-tailed Student’s t-test.

## Discussion

In this study, we observed that the upregulation of LMO3 is associated with the differentiation into the classical/progenitor subtype and predicts a favorable prognosis for patients with PDAC. Elevated LMO3 levels induced changes in cancer cell metabolism, resulting in inhibited proliferation and reduced migration and invasion in PDAC cells. The presence of LMO3-induced G3P was correlated with increased patient survival. Additionally, the upregulation of LMO3 led to a decrease in the basal-like/squamous gene signature in both patient PDAC samples and PDAC cells.

Lim domain only (LMO) proteins constitute a family of transcription co-factors recognized for their roles in cell differentiation and development^32, 33^. This family comprises four members: LMO1, LMO2, LMO3, and LMO4^32, 33^. Among these, LMO1 and LMO2 have been identified as oncogenes in T-cell leukemia^34–36^, while LMO4 overexpression promotes breast cancer development^37, 38^. LMO3, primarily expressed in the brain^33^, plays a role in the development of neuroblastoma through promoting HEN2 activity^39^ and downregulating p53-dependent mRNA expression^40^. It has also been linked to the aggressiveness of hepatocellular carcinoma^41^ and lung cancer^42^. However, the function of LMO3 in cancers remains controversial. Previous studies in PDAC and prostate cancer have suggested that LMO3 may exert suppressive effects on tumorigenesis in these cancers^43, 44^. Furthermore, LMO3 may be involved in metabolic reprogramming^45^ and has been identified as a master regulator of adipogenesis^24^. Nevertheless, the role of LMO3 in PDAC has not been fully elucidated. Our results reveal that LMO3 functions as an inducer of classical/progenitor differentiation, leading to reduced cellular activity and improved patient survival.

Cancer cells undergo metabolic reprogramming characterized by increased glycolysis, lipid synthesis, and amino acid production, often involving the pentose phosphate pathway^8, 9^. Previous research has shown that the basal-like/squamous subtype of cancer relies on heightened glycolysis and amino acid metabolism, while the classical/progenitor subtype is associated with elevated lipogenesis^13–17^. Amino acids play a crucial role in PDAC development^13, 46, 47^, while the effects of lipids are more intricate^25, 47, 48^. For instance, previous studies, including our own, have indicated that fatty acids can inhibit the growth of PDAC^25–27^. Consistently, LMO3-induced metabolic changes may promote the classical/progenitor subtype by downregulating amino acid metabolism while increasing lipogenesis. Conversely, its loss may redirect differentiation toward the basal-like/squamous subtype, characterized by increased glycolysis and reduced lipogenesis. In our *in silico* and *in vitro* studies, we identified six metabolites commonly upregulated in LMO3-high PDAC tumors and PDAC cells, classified as lipids. Notably, glycerol-3-phosphate (G3P) emerged as one of the most significantly upregulated metabolites in LMO3-high PDAC and the classical/progenitor subtype. G3P has been found to exert inhibitory effects on tumorigenesis in various cancers, including prostate cancer, lung cancer, colorectal cancer, and breast cancer^31^. The increase in G3P levels in the classical/progenitor subtype PDAC tumors is likely mediated by GPD1, an enzyme responsible for synthesizing G3P from DHAP. In agreement with the inverse relationship between G3P level and basal-like/squamous subtype differentiation, we observed that high levels of G3P in PDAC are associated with increased patient survival, and the upregulation of G3P-producing GPD1 is evident in LMO3-high PDAC and non-basal-like/squamous subtype tumors.

A limitation of this study originates from the absence of mechanistic analyses concerning the interaction between LMO3 and other proteins, such as HEN2^39^ and p53^40^. While the upregulation of LMO3 resulted in lipogenesis, reduced cellular activity, induction of the classical/progenitor subtype, and extended patient survival in PDAC, it is premature to label LMO3 as a suppressor of PDAC. It remains possible that LMO3 might act as an oncogene related to classical/progenitor subtype differentiation through interactions with other proteins. The observed extension in patient survival could be attributed to the induction of classical/progenitor differentiation, a subtype associated with favorable survival due to increased lipogenesis^26, 27^. Furthermore, this study focused on gain-of-function experiments because all the PDAC cell lines we utilized exhibited downregulated LMO3 expression. Future investigations should include mechanistic studies and loss-of-function experiments to provide a more comprehensive understanding of LMO3’s role in PDAC.

In summary, our findings indicate that LMO3 expression promotes transcriptomic and metabolomic features of the classical/progenitor subtype, while its downregulation supports differentiation into basal-like/squamous tumors and increased PDAC aggressiveness. Furthermore, LMO3 promotes a distinct metabolic profile characterized by heightened lipogenesis and reduced amino acid metabolism, a hallmark of the classical/progenitor PDAC subtype.

## Author Contributions

**Yuuki Ohara:** Conceptualization, Methodology, Resources, Investigation, Writing – Original Draft, Writing – Review & Editing, Project Administration. **Amanda J. Craig**: Methodology, Data Curation, Writing – Review & Editing, Project Administration. **Huaitian Liu:** Data Curation, Writing – Review & Editing, Project Administration. **Shouhui Yang:** Methodology, Writing – Review & Editing, Project Administration. **Paloma Moreno:** Project Administration. **Tiffany H. Dorsey:** Project Administration. **Helen Cawley:** Project Administration. **Azadeh Azizian:** Resources, Project Administration. **Jochen Gaedcke:** Resources, Project Administration. **Michael Ghadimi:** Resources, Project Administration. **Nader Hanna:** Resources, Project Administration. **Stefan Ambs:** Methodology, Writing – Review & Editing, Supervision. **S. Perwez Hussain:** Conceptualization, Writing – Review & Editing, Supervision, Funding Acquisition. The work reported in the paper has been performed by the authors, unless clearly specified in the text.

## Acknowledgments

The authors express their gratitude to the dedicated staff and study coordinators at the University of Maryland School of Medicine for their valuable assistance in procuring clinical biospecimens and patient data. Additionally, we extend our appreciation to the Department of General, Visceral, and Pediatric Surgery at the University Medical Center Göttingen, Göttingen, Germany, for their generous support in handling clinical samples. Finally, special thanks to the dedicated team at the NCI-CCR Sequencing Facility Frederick for their invaluable contributions to this research.

## Data and materials availability

The data that support the findings of this study are available from the corresponding author upon request.

## Ethics Statement

Pancreatic tissues were obtained from surgically resected PDAC patients at the University of Maryland Medical System (UMMS) in Baltimore, MD, under the purview of an NCI-UMD resource contract, as well as from the University Medical Center Göttingen, Germany. Board-certified pathologists evaluated PDAC histopathology. The utilization of these clinical specimens in our research was subject to review by the NCI-Office of the Human Subject Research Protection (OHSRP) at the NIH in Bethesda, MD (Exempt#4678). Informed written consent was obtained from all participants, and all procedures were conducted in strict adherence to ethical standards and in accordance with the 1975 Declaration of Helsinki, as revised in 2008.

## Abbreviations

ADEX: aberrantly differentiated endocrine exocrine
DAB - 3: 3’-diaminobenzidine
DHAP: dihydroxyacetone phosphate
G3P: glycerol 3-phosphate
GPD: glycerol-3-phosphate dehydrogenase
GSEA: Gene Set Enrichment Analysis
IHC: immunohistochemistry
IPA: Ingenuity pathway analysis
LMO: LIM Domain Only
PDAC: pancreatic ductal adenocarcinoma
STR: short tandem repeat

## Supplementary figure legends

**Supplementary Figure 1.**
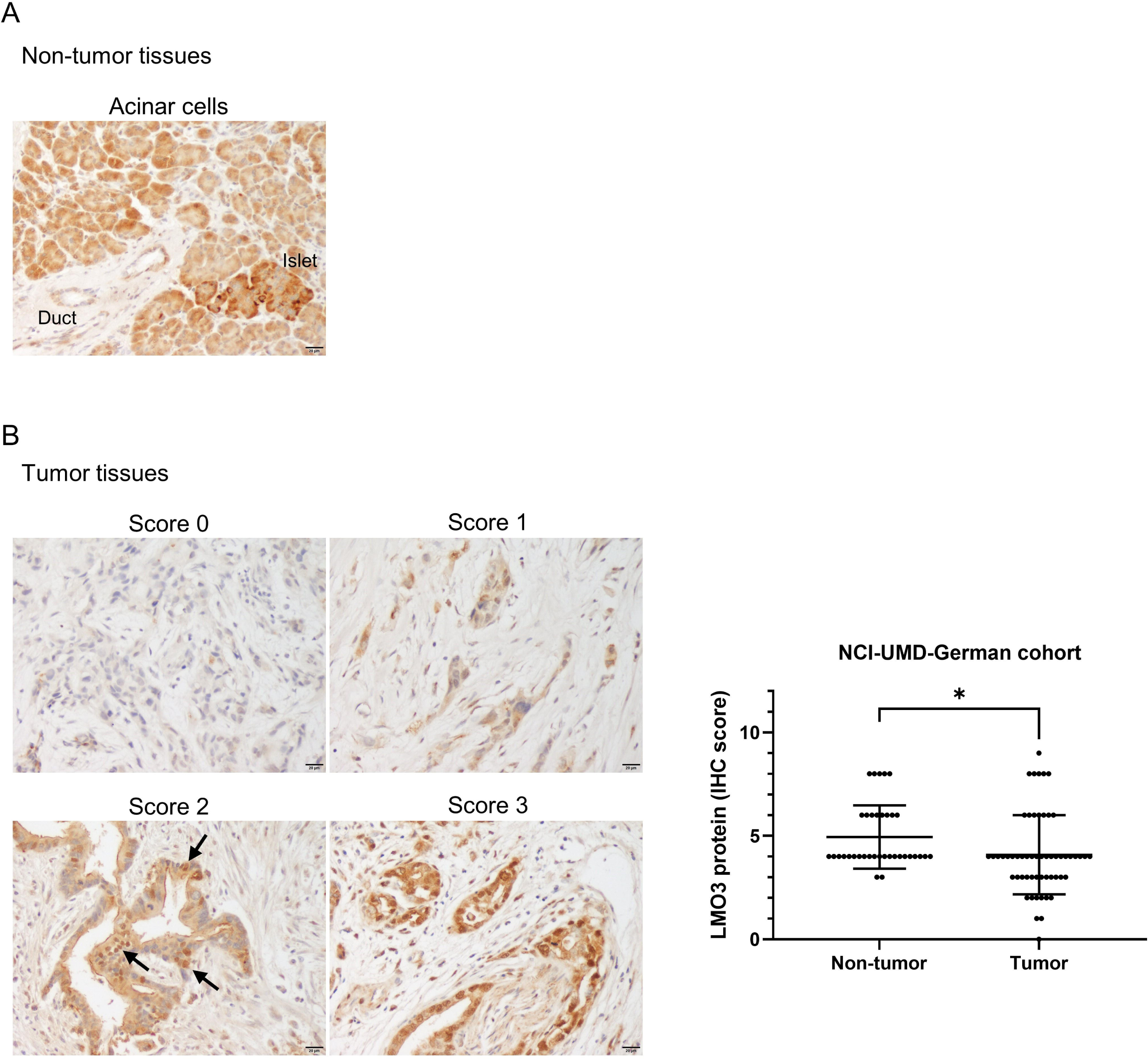
Downregulation of LMO3 protein in PDAC tumors in the NCI-UMD-German cohort. IHC of LMO3 in tumor and non-tumor tissue sections of PDAC patients. LMO3 protein is detected in the cytoplasm, as shown by the brown DAB-based IHC in the tumor cells, the acinar cells, and the endocrine cells in pancreatic islets. The protein is also detected in the nucleus of tumor cells. (A) Representative LMO3 protein staining in non-cancerous acinar cells (intensity score 2 in acinar cells and score 3 in islets). (B) LMO3 in representative tumor sections (scores 0-3), indicating downregulation of LMO3 in tumor cells when compared to the non-tumor acinar cells (see graph to the right). The staining strength is categorized as follows: score 0 (non-stained), score 1 (weak), score 2 (moderate), and score 3 (strong). The presence of the protein is detected in the nucleus of tumor cells (arrows) as well as in the cytoplasm. Scale bar is 20 μm. More details can be found in Materials and Methods. Data represent mean ± SD. *p ≤ 0.05, **p ≤ 0.01, ***p ≤ 0.001, ****p ≤ 0.0001 by unpaired two-tailed Student’s t-test. DAB; 3, 3’-diaminobenzidine, IHC; immunohistochemistry.

**Supplementary Figure 2.**
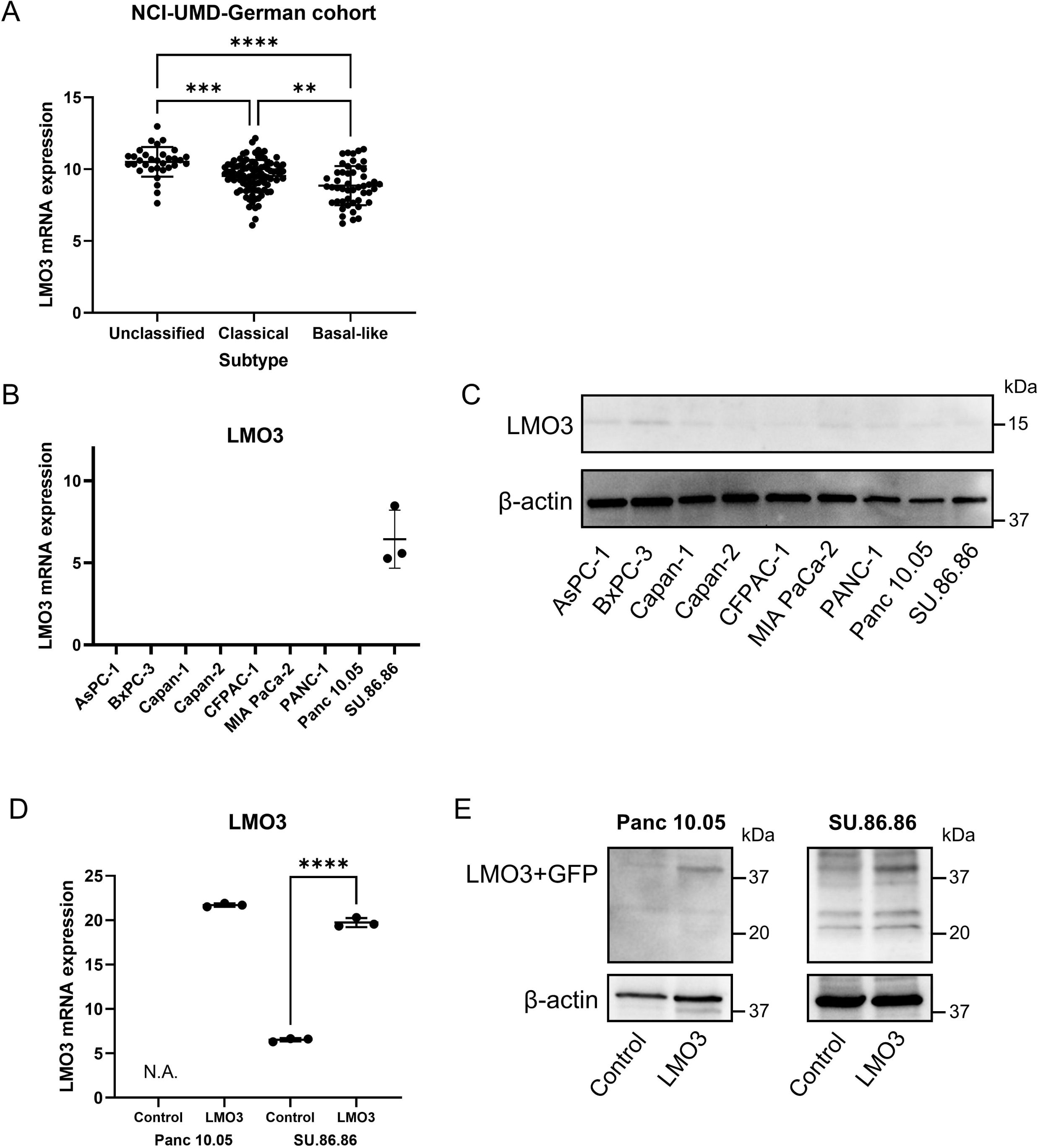
LMO3 expression in human PDAC cells. (A) *LMO3* mRNA expression by subtype in the NCI-UMD-German cohort. (B-C) Endogenous levels of LMO3 mRNA and protein in various human PDAC cell lines, revealing consistent suppression of LMO3 expression across all examined PDAC cell lines. (D-E) Confirmation of LMO3 transgene overexpression at the mRNA (bar graph) and protein levels in Panc 10.05 and SU.86.86 cells. Data represent mean ± SD. *p ≤ 0.05, **p ≤ 0.01, ***p ≤ 0.001, ****p ≤ 0.0001 by ANOVA (A) or unpaired two-tailed Student’s t-test (D).

**Supplementary Figure 3.**
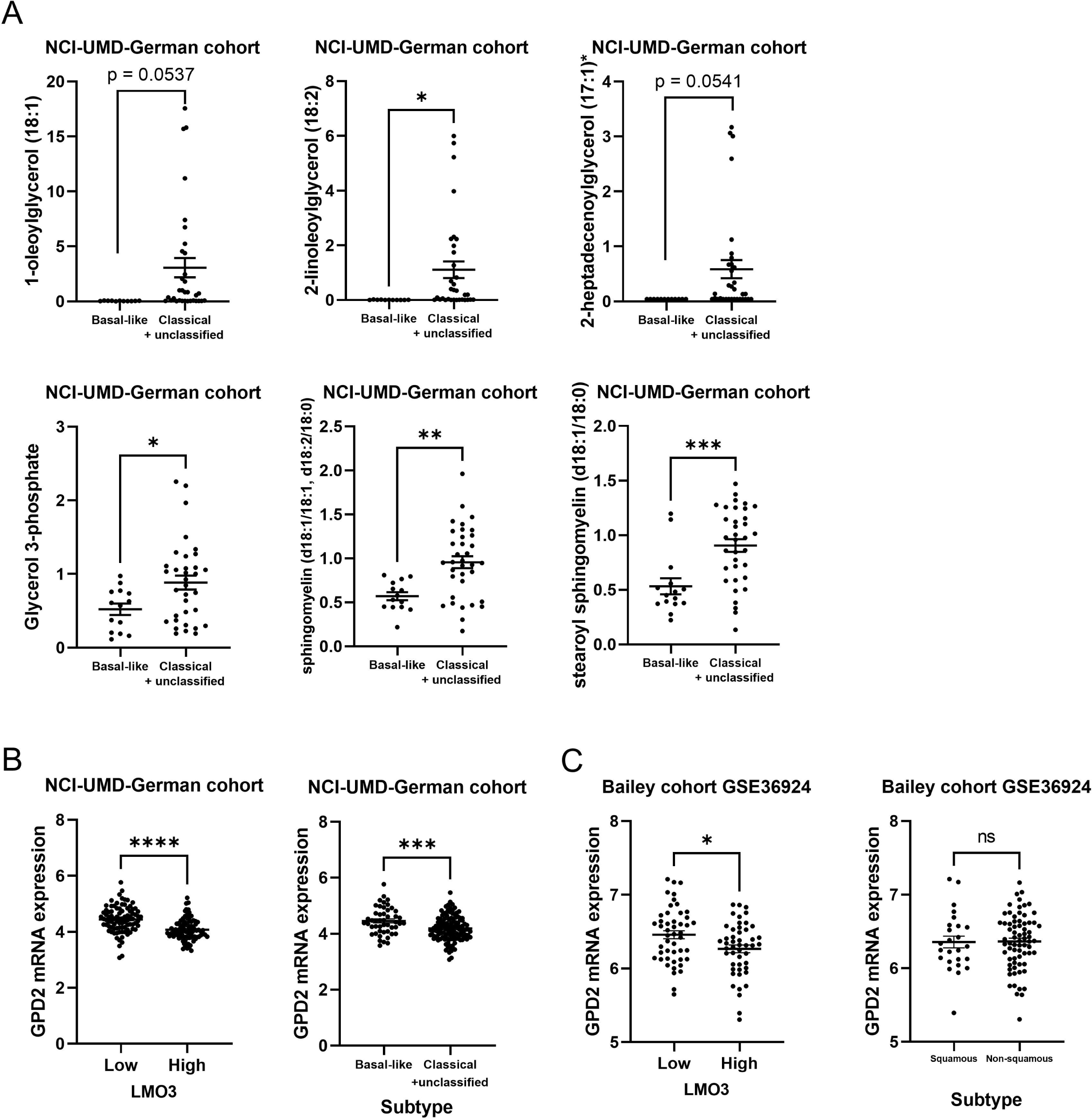
Upregulated metabolites and GPD2 expression in PDAC cohorts. (A) Upregulated metabolites in LMO3-high patient PDAC tumors and in LMO3-overexpressing Panc 10.05 cells (Figure 5A) were also downregulated in the non-basal-like/squamous PDAC (non-basal-like/squamous: classical/progenitor + unclassified PDAC; n = 35, basal-like/squamous PDAC; n = 15). GraphPad analysis was used to exclude the outliers from the graphs. (B) *GPD2* was downregulated both in LMO3-high PDAC and non-basal-like/squamous subtype (classical/progenitor + unclassified PDAC) in the NCI-UMD-German cohort. (C) *GPD2* was downregulated in LMO3-high PDAC in the Bailey cohort^3^. Data represent mean ± SD. *p ≤ 0.05, **p ≤ 0.01, ***p ≤ 0.001, ****p ≤ 0.0001 by unpaired two-tailed Student’s t-test.

